# Assessment of Microbial Consortia for Biogenic Mineralization Potential of Lead (Pb) using Microbiologically Induced Calcite Precipitation and Nitrate Reduction

**DOI:** 10.64898/2026.09.22.753425

**Authors:** Ayana Bhattacharya, Sneha Hiremath, Deepesh Nagarajan, Tanushree Ghosh

## Abstract

Groundwater contamination by lead (Pb^+2^) and nitrate (NO₃⁻) exceeding the permissible limits of the World Health Organization (WHO) and the Bureau of Indian Standards (BIS) (10500:2012) of 10 ppb Pb^+2^ and 45–50 ppm NO₃⁻, poses a severe health risk of systemic toxicity in humans and animals. Microbial nitrate reduction coupled with microbiologically induced calcite precipitation (MICP) proved to be a novel bioremediation strategy to transform soluble lead nitrate [Pb(NO₃)₂] into insoluble lead biominerals. This study presents statistical optimization of operational parameters for nitrate reduction and co-precipitation of biogenic lead carbonate-calcium carbonate by nitrate-reducing bacterial consortia. The SEM, TEM with SEAD, and XRD analysis of the biomineral confirmed PbCO₃ co-precipitaion with CaCO_3_. A two-level factorial design was applied to screen significant variables and interactive factors that influence nitrate reduction and biomineral precipitation. The highest nitrate reduction, maximum ammonia production, and biogenic Pb-mineralization efficiency of> 90% were obtained at the optimum conditions (initial nitrate requirement of 600ppm, temperature 37 °C, pH 8, and carbon content of 5.5 g/L) compared to those obtained under the common process parameters. ANOVA with a high correlation coefficient (R^2^ > 0.99) corroborated the linear model for the biogenic mineralization process. This study on the biogenic removal of Pb by nitrate-reducing bacterial consortia revealed possible opportunities for large-scale treatment of Pb-contaminated surface and sub-surface water bodies.

## 1. INTRODUCTION

Lead (Pb^+2^) contamination in groundwater is posing a systemic toxicity threat to human and environmental health. Groundwater contributes to about 97% of global freshwater and serves as the single largest supply of global drinking water [1]. Anthropogenic and geogenic contributions are the major routes of lead in groundwater. The widespread occurrence of Pb^+2^ in the environment is primarily attributed to anthropogenic sources, while other sources can be geogenic [2]. The World Health Organization (WHO) has established a permissible limit of 10 µg L⁻¹ (ppb) for lead in potable water and also reported an estimated 143,000 deaths per year attributable to lead poisoning, accounting for about 0.6% of the global disease burden [3]. Using the 2019 Global Burden of Disease dataset, approximately 815 million children worldwide are estimated to have blood lead levels (BLL) above 5 µg/dL, with nearly 50% of these children residing in Southeast Asia [4]. The global scenarios of Pb-contaminated hotspots reported > 1000 ppb of Pb include Bangladesh (shallow tube wells up to 1,167 µg/L) [5], Indonesia’s Maros (rainy-season groundwater up to 9,280 µg/L) [2], Nigeria’s Abakaliki (mining groundwater up to 38,000 µg/L) [5], and India’s Punjab (the Malwa region up to 28,040 µg/L) [6]. The Institute for Health Metrics and Evaluation found that India itself experienced nearly 7 million lead-attributable Disability-Adjusted Life Years (DALY) and more than 2.3 lakh deaths in 2019 [7]. The alluvial aquifer system of northwest India records elevated lead (>10 µg/L) content in shallow urban groundwater, with the contamination traced to urbanization and industrial discharge. In southern India, the Kabini River basin in the Western Ghats shows lead above the permissible limit in the majority of sampled groundwater: monsoon concentrations span 0.01–0.98 mg/L (mean 0.4 mg/L) and non-monsoon values 0.01–0.17 mg/L (mean 0.10 mg/L), attributed by the authors to industrial effluents and agricultural inputs [8]. The aqueous chemistry of lead indicates that lead nitrate, Pb(NO₃)₂, is highly soluble (59.7 g/100 g H₂O at 25°C) [9], whereas oxides, hydroxides, and carbonates of lead are sparingly soluble. Elevated nitrate levels in the groundwater are also increasing lead solubility [9]. Lead dissolution and mobility are highly affected by the pH and available nitrate. In a classic experimental phase study of the Pb²⁺–NO₃⁻–H₂O system, Grimes and colleagues reported that raising the pH in aqueous lead nitrate solutions first precipitates the tetranuclear cluster [Pb₄₄][NO₃]₄; above pH ∼5.25, nitrate in the lattice is progressively replaced by hydroxide, yielding distinct phases at pH ∼7 ([Pb₃₄]²⁺-based) and ∼8.5 ([Pb₆O₆]⁴⁺-based) [10]. This study indicated that the presence of nitrate highly influences lead transport in the aqueous environment. Nitrate stabilizes soluble/hydroxo-cluster species at low pH, while nitrate replacement by OH⁻ at higher pH drives precipitation. The reversal mechanism by denitrification also proved to be a recommended solution to combat groundwater lead load. Nitrate consumption by dissimilatory reduction raises pH, facilitates carbonate precipitation, and removes Pb as Pb⁰, PbS, PbO, PbSO₄, and pyromorphite at up to 99% efficiency [11–14].

Microbial remediation is emerging as a cost-effective, efficient, and eco-friendly alternative to traditional physical or chemical methods, operating through biosorption, bioprecipitation, biomineralization, and bioaccumulation, and through bacterial generation of nanoparticles that convert dissolved Pb into insoluble removable forms [15]. Among the microbial strategies, dissimilatory nitrate reduction is mechanistically central: nitrate acts as a terminal electron acceptor through the enzyme dissimilatory nitrate reductase, with nitrate usage most prominent at lower Pb concentrations (80–250 ppm) [14]. Bacterial nitrate dissimilation also drives microbially induced carbonate precipitation (MICP), with a highly negative standard Gibbs free energy (ΔG° = 785 kJ/mol) and approximately two-fold higher carbonate yield than ureolysis, simultaneously addressing nitrate pollution and metal detoxification [11]. Continuous anoxic bioreactor systems have demonstrated up to 99% Pb removal efficiency with denitrification as the dominant detoxification mechanism, supported by biosorption, sulfur-reducing bacterial activity, bioprecipitation, and bioremoval [12]. These findings position nitrate-reduction-coupled lead bioremediation as a viable, sustainable technology for contaminated sites such as urban shallow aquifers, smelter-affected zones, and unregulated battery-recycling areas [12,15]. Microbially induced carbonate precipitation (MICP) has emerged as a pioneering, cost-effective, and environmentally compatible alternative that converts soluble, bioavailable lead into sparingly soluble carbonate minerals, permanently sequestering the contaminant [16,17]. MICP exploits urease-producing microorganisms that hydrolyze urea to raise the pH and generate carbonate ions, which precipitate dissolved metals, including Pb²⁺ as insoluble carbonate phases such as cerussite (PbCO₃) and/or hydroxyserussite ((Pb₃(CO₃)₂(OH)₂) or entrap them within the lattice of calcium carbonate (calcite) crystals [16,17]. Because the process is independent of metal valence state, toxicity, and redox potential, ureolytic biomineralization is regarded as one of the most robust biological strategies for lead immobilization in both aqueous and soil systems [17].

The MICP mechanism has been reported in our previous study in agreement with other well-studied literature [16–19] and proceeds as follows (Eq 1-7) :

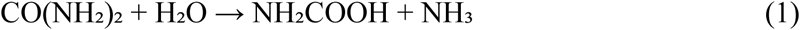

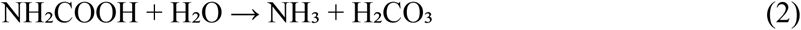

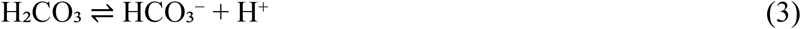

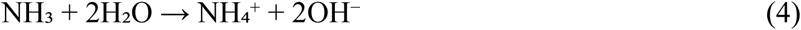

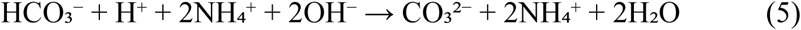

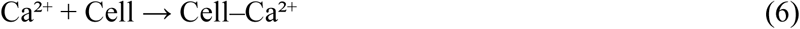

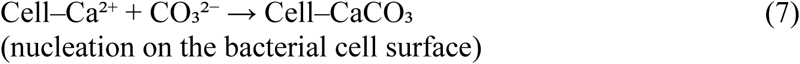

The hydrolysis products equilibrate in water to form bicarbonate, ammonium, and hydroxyl ions. The phenomenon leads to elevated pH and shifts the bicarbonate equilibrium toward carbonate ion formation. The metabolic CO₂ from respiration further increases dissolved inorganic carbon in the microenvironment, which in turn enhances precipitation [17]. High pH favours CO₃²⁻ formation from HCO₃⁻ [17], and in a urea-hydrolysis system the NH₄⁺/NH₃ and HCO₃⁻/CO₃²⁻ equilibria converge at a pH range of 8.5-9.3 [20]. Reported enzyme kinetic parameters illustrate the biochemical basis of the process: for *Sporosarcina pasteurii*, urease activity decreased at pH 7.0 with Km and Vmax values of 41.6 mM and 3.55 mM min⁻¹ mg⁻¹ of protein, respectively [18]. The enzyme urease contains two nickel ions at the centre of the protein. At the structural level, urease ligates urea to a dinuclear nickel centre; urea binds Ni-1 to complete its tetrahedral coordination while one hydroxide ligand of Ni-2 attacks the carbonyl carbon and releases NH₄⁺ and CO₂/CO₃²⁻, and elevates the pH from 7.5 to 9.1 [21]. The calcite-depositing capacity of ureolytic bacteria leads to mineralization of soluble heavy metal ions and their ultimate conversion to carbonates, with CaCO₃ representing a stable mineral phase that provides permanent sequestration [22]. The geochemical rationale for the MICP approach to lead is that carbonates play a major role in lead speciation in aqueous environments by forming insoluble compounds [2]. At pH ≥ 6.6, PbCO₃ (cerussite) can precipitate [23], and ureolytic bacteria that hydrolyse urea efficiently generate carbonate ions and elevate the pH to alkaline conditions (8.0–9.1), which promotes the precipitation of lead and calcium carbonate [23]. The Pb-related precipitate chemistry is dominated by competition between Pb²⁺ and Ca²⁺ for carbonate: scanning electron microscopy–energy-dispersive X-ray spectroscopy (SEM– EDS) studies have shown that cerussite precipitates before calcite because Pb²⁺ has higher affinity for CO₃²⁻ and OH⁻ than Ca²⁺ [24]. Consequently, in mixed systems, lead is either precipitated directly as PbCO₃ or coprecipitated and entrapped within calcite lattices, limiting subsequent metal leaching [11]. There are many bacterial strains reported to facilitate Pb-biomineralization through the urolytic pathway (Table 1), while only a few combined the nitrate reduction with the MICP. A well recognized limitation of MICP for lead is that Pb-related precipitates can potentially leach under harsh environmental conditions, releasing acid and threatening human health. Self-healing microbial-induced calcium carbonate materials within which spores germinate, transform to vegetative cells, secrete urease, and hydrolyze urea to re-precipitate carbonate after leaching episodes. This can maintain an immobilization efficiency greater than 95% after five harsh-environment cycles, whereas non-self-healing controls decayed to less than 10% after three cycles [24]. Spores and vegetative cells provided extra nucleation sites for Pb²⁺ and minerals, and extracellular polymeric substances bound Pb²⁺ through functional groups and chemical bonds, preventing migration [24]. Entrapment of PbCO₃ within the calcite lattice similarly limited leaching of metals from the carbonate-bound complex to the surrounding environment [11]. These data indicate that MICP-based lead carbonate precipitation can achieve the long-term stability required of a remediation endpoint, provided the biological carbonate-replenishment capacity is retained.

**Table 1:**
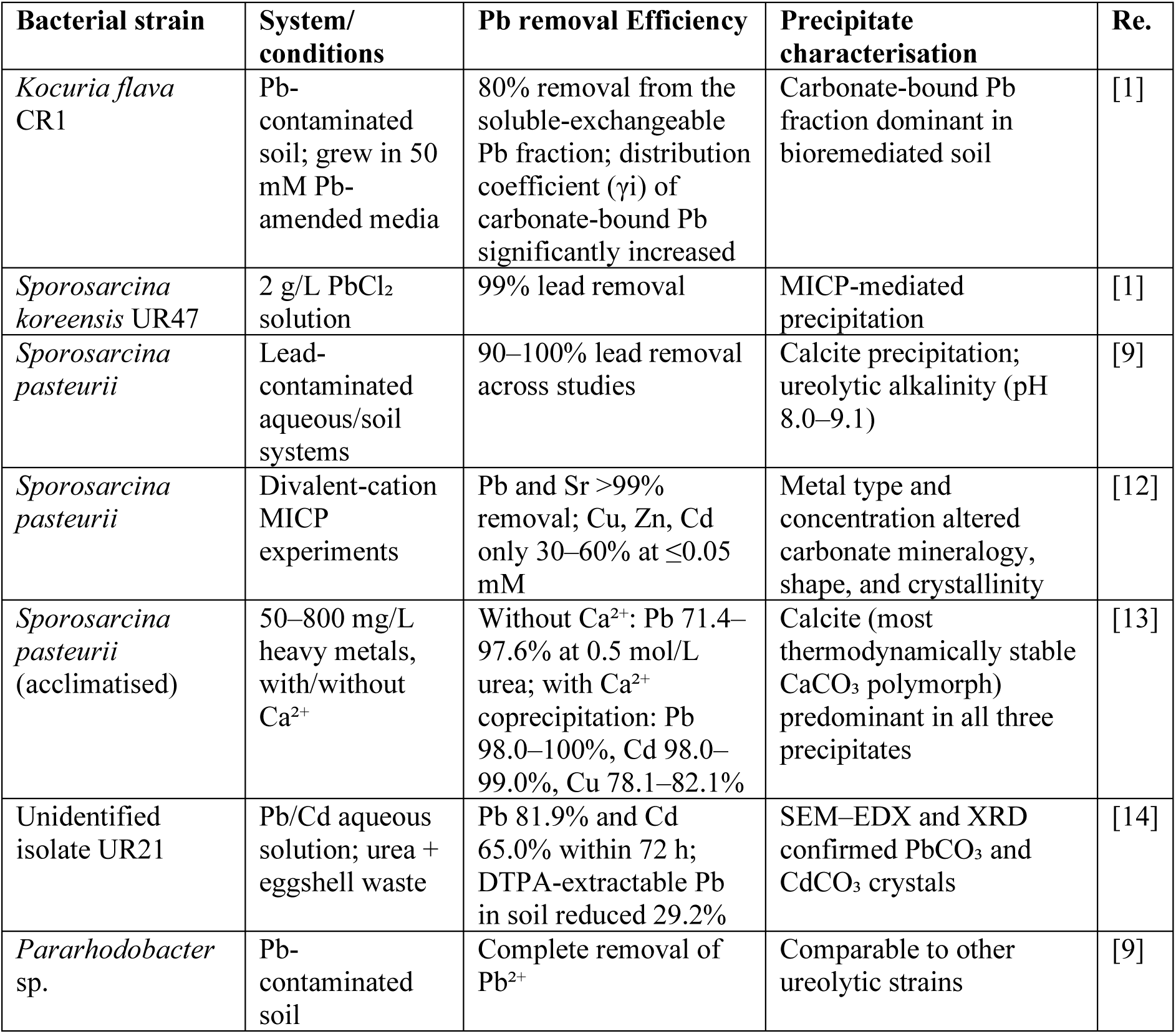
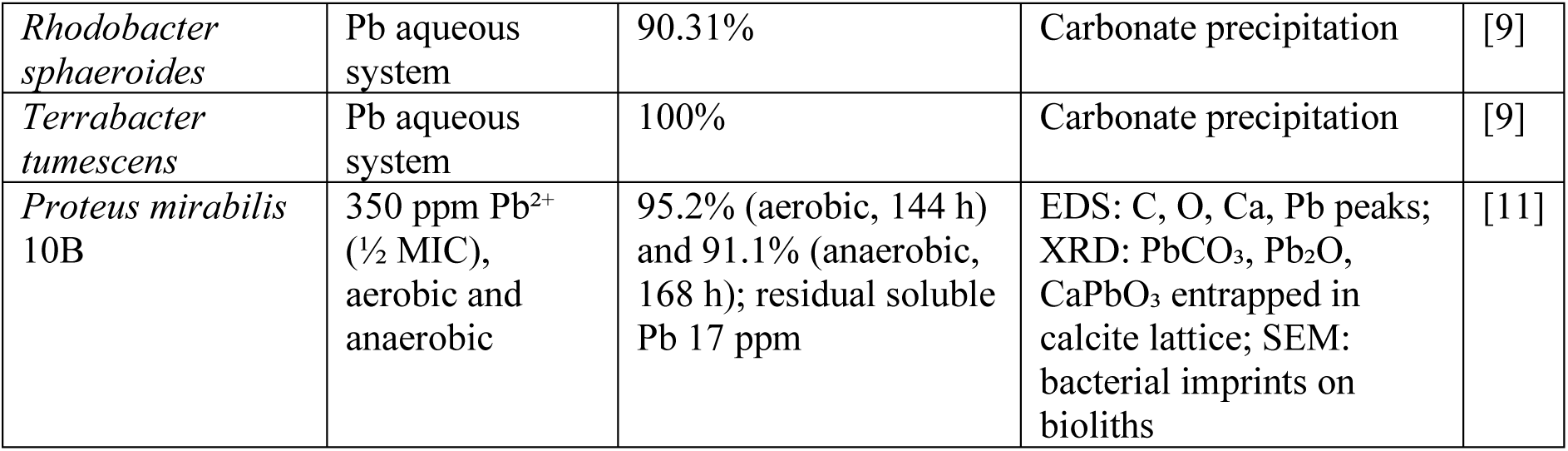
Summary of bacterial strains involved in biogenic Pb precipitation.

| Bacterial strain | System/<br>conditions | Pb removal Efficiency | Precipitate<br>characterisation | Re. |
| --- | --- | --- | --- | --- |
| *Kocuria flava*<br>CR1 | Pb-contaminated<br>soil; grew in 50<br>mM Pb-<br>amended media | 80% removal from the<br>soluble-exchangeable<br>Pb fraction; distribution<br>coefficient ($\gamma_i$) of<br>carbonate-bound Pb<br>significantly increased | Carbonate-bound Pb<br>fraction dominant in<br>bioremediated soil | [1] |
| *Sporosarcina*<br>*koreensis* UR47 | 2 g/L $PbCl_2$<br>solution | 99% lead removal | MICP-mediated<br>precipitation | [1] |
| *Sporosarcina*<br>*pasteurii* | Lead-<br>contaminated<br>aqueous/soil<br>systems | 90–100% lead removal<br>across studies | Calcite precipitation;<br>ureolytic alkalinity (pH<br>8.0–9.1) | [9] |
| *Sporosarcina*<br>*pasteurii* | Divalent-cation<br>MICP<br>experiments | Pb and Sr >99%<br>removal; Cu, Zn, Cd<br>only 30–60% at $\leq 0.05$<br>mM | Metal type and<br>concentration altered<br>carbonate mineralogy,<br>shape, and crystallinity | [12] |
| *Sporosarcina*<br>*pasteurii*<br>(acclimatised) | 50–800 mg/L<br>heavy metals,<br>with/without<br>$Ca^{2+}$ | Without $Ca^{2+}$: Pb 71.4–<br>97.6% at 0.5 mol/L<br>urea; with $Ca^{2+}$<br>coprecipitation: Pb<br>98.0–100%, Cd 98.0–<br>99.0%, Cu 78.1–82.1% | Calcite (most<br>thermodynamically stable<br>$CaCO_3$ polymorph)<br>predominant in all three<br>precipitates | [13] |
| Unidentified<br>isolate UR21 | Pb/Cd aqueous<br>solution; urea +<br>eggshell waste | Pb 81.9% and Cd<br>65.0% within 72 h;<br>DTPA-extractable Pb<br>in soil reduced 29.2% | SEM–EDX and XRD<br>confirmed $PbCO_3$ and<br>$CdCO_3$ crystals | [14] |
| *Pararhodobacter*<br>sp. | Pb-contaminated<br>soil | Complete removal of<br>$Pb^{2+}$ | Comparable to other<br>ureolytic strains | [9] |
| *Rhodobacter sphaeroides* | Pb aqueous system | 90.31% | Carbonate precipitation | [9] |
| *Terrabacter tumescens* | Pb aqueous system | 100% | Carbonate precipitation | [9] |
| *Proteus mirabilis* 10B | 350 ppm Pb<sup>2+</sup> (½ MIC), aerobic and anaerobic | 95.2% (aerobic, 144 h) and 91.1% (anaerobic, 168 h); residual soluble Pb 17 ppm | EDS: C, O, Ca, Pb peaks; XRD: PbCO<sub>3</sub>, Pb<sub>2</sub>O, CaPbO<sub>3</sub> entrapped in calcite lattice; SEM: bacterial imprints on bioliths | [11] |

Many physicochemical factors influence lead carbonate precipitation via MICP. The most studied factors are urea and calcium concentration. In the absence of Ca²⁺, increasing urea concentration enhances carbonate supply to improve Cd²⁺ removal to 89.9–99.7% at 2.0 mol/L urea, whereas Pb²⁺ removal was optimal at a lower urea concentration of 0.5 mol/L (efficiency of 71.4–97.6%). Concentrations increasing beyond 0.5 mol/L of urea showed a little positive effect on lead removal [25]. However, coprecipitation of Pb²⁺ with Ca²⁺ dramatically improves lead removal. In the presence of Ca²⁺, Pb removal peaks to 98.0–100%, while it remains in the range 71.4–97.6% without Ca²⁺, because Pb²⁺ substitutes into the growing calcite lattice [25]. Elevated Ca²⁺ levels in groundwater modulate overall MICP performance [16]. Another important factor is the pH. Ureolytic activity raises pH to the alkaline window (8.0–9.1) at which both PbCO₃ and CaCO₃ precipitate [23]; the HCO₃⁻/CO₃²⁻ equilibrium converges near pH 9.3 [20], and the urease-mediated pH rise (7.5 to 9.1) is the rate-limiting driver of the carbonate supersaturation [21].

Nitrate-reduction-driven carbonate precipitation is a mechanistically distinct but converging route to bacterial lead carbonate precipitation. In a study with *P. mirabilis* 10B cultures, nitrate reductase activity showed a positive correlation with bacterial growth, pH, NO₃⁻ and NO₂⁻ consumption, and removal of Ca²⁺, Pb²⁺, and Hg²⁺ [11]. Denitrification resulted in elevated pH to 9.3 in the biotic control and to 8.9 in remediated samples due to ammonia production. This leads to the precipitation of 1200, 1008, and 925 ppm of Ca²⁺ as CaCO₃ crystals with 100%, 84%, and 77% removal efficiency, respectively. This also influenced anoxic precipitation of Pb²⁺ by about 91.1% [11]. Aerobic removal (95.2% for Pb²⁺) was reported to be faster and slightly higher, attributed to the higher redox potential supporting greater nitrate reduction, better bacterial metabolic activity, and proliferation resulting in increased nucleation sites [11,19]. The denitrification-driven carbonate precipitation has not been explored in depth, while there is very little reporting on the parametric optimization to maximize Pb²⁺-biomineralization.

The present study focused on the assessment of microbial consortia for nitrate reduction and ureolytic capability. The nitrite and ammonia production rates and biomineral precipitation ability were compared to screen the different BW samples in the presence and absence of Pb²⁺. The best-responding GW8 consortia was selected for the parametric optimization using a two-level factorial to maximize Pb²⁺-biomineral precipitation. This study explored the possibility of lab-scale to pilot-scale translation of biogenic Pb²⁺ remediation and recovery.

## 2. MATERIALS AND METHODS

### 2.1. Materials

All chemicals and reagents used in this study were of analytical grade and used without further purification unless otherwise stated. Sulphanilic acid solution (Griess Reagent A, 0.8%, containing sulphanilic acid in acetic acid, Cat. No. R015) and α-naphthylamine solution (Griess Reagent B, Cat. No. R009), Sodium nitrite (NaNO₂, 98.0%, Cat. No. GRM417), Ammonium chloride (NH₄Cl, ≥99.0%, Cat. No. GRM717), Rochelle salt solution (potassium sodium tartrate) and Nessler’s reagent (mixture, LR grade, Cat. No. REA185), Potassium nitrate (KNO₃, 99.00–100.50%, Cat. No. GRM3946), Potassium chloride (KCl, 98.00–102.00%, Cat. No. GRM697), Potassium dihydrogen phosphate (KH₂PO₄), Magnesium sulphate heptahydrate (MgSO₄·7H₂O, 99.00–102.00%) Cat. No. GRM683), Ferrous sulphate heptahydrate (FeSO₄·7H₂O, 98.00–104.50% Cat. No. GRM372), Urea [CO(NH_2_)_2_], 99.00–100.50%, Cat.

No. GRM3976), Calcium chloride (anhydrousCaCl₂, 93.00–102.00%, Cat. No. GRM710), D-(+)-glucose anhydrous (C_6_H_12_O_6_), and Lead (II) nitrate [Pb(NO_3_)_2_], ≥99.0%, Cat. No. GRM3920) were purchased from HiMedia Laboratories Pvt Ltd., Mumbai, India. Nitrate broth (NB), nitrate agar (NA), and Luria–Bertani (LB) broth and a modified Nitrate Salt Medium (NSM) were used for culturing and storage of microbial strains. The NB, NA, and LB were purchased from HiMedia Laboratories Pvt Ltd., Mumbai, India, and were prepared according to the manufacturer’s instructions. The nitrate broth supplemented with urea (20 mg L⁻¹)) and CaCl₂ (1 mg L⁻¹) is referred to as NBUC medium and the composition can be found in our previous study [19].

The NSM basal medium was prepared using KCl (1.0 gL⁻¹), KH₂PO₄ (0.5 gL⁻¹), MgSO₄·H₂O (0.5 gL⁻¹) ), FeSO₄ (1 mgL⁻¹), KNO₃ (1.0 gL⁻¹), CaCl₂ (1 mg L⁻¹), Urea (20gL⁻¹) and Glucose (2.75gL⁻¹) in MilliQ water. The first five media components were autoclaved (121 °C and 15 psi) separately in the 800 mL basal stock, whereas CaCl_2_, urea and glucose were filter-sterilized separately in the remaining 200 mL to prepare NSM complete media. Sterile Pb(NO₃)₂ to a final Pb²⁺ concentration of 100 µg L⁻¹ (100 ppb) was used for all the Pb²⁺ biomineralization experimental conditions.

### 2.2. Sample collection and Microorganism

Twelve groundwater samples were collected from Chintamani village, Chikballapura District, Karnataka, India for this study. All borewell (BW) water samples were collected using the purge and sample method. Each borewell was pumped for five minutes to remove stagnant water from the well casing and tubing. After purging, 2 L water samples were collected in polypropylene bottles. All the polypropylene bottles were pre-contaminated with the water samples to be collected to avoid cross-contamination. Figure 1 shows the geographical locations of the water collection sites. The latitude-longitude of sampling sites, chemical and elemental parameters of the BW water samples were reported in our previous study [28].

**Figure 1:**
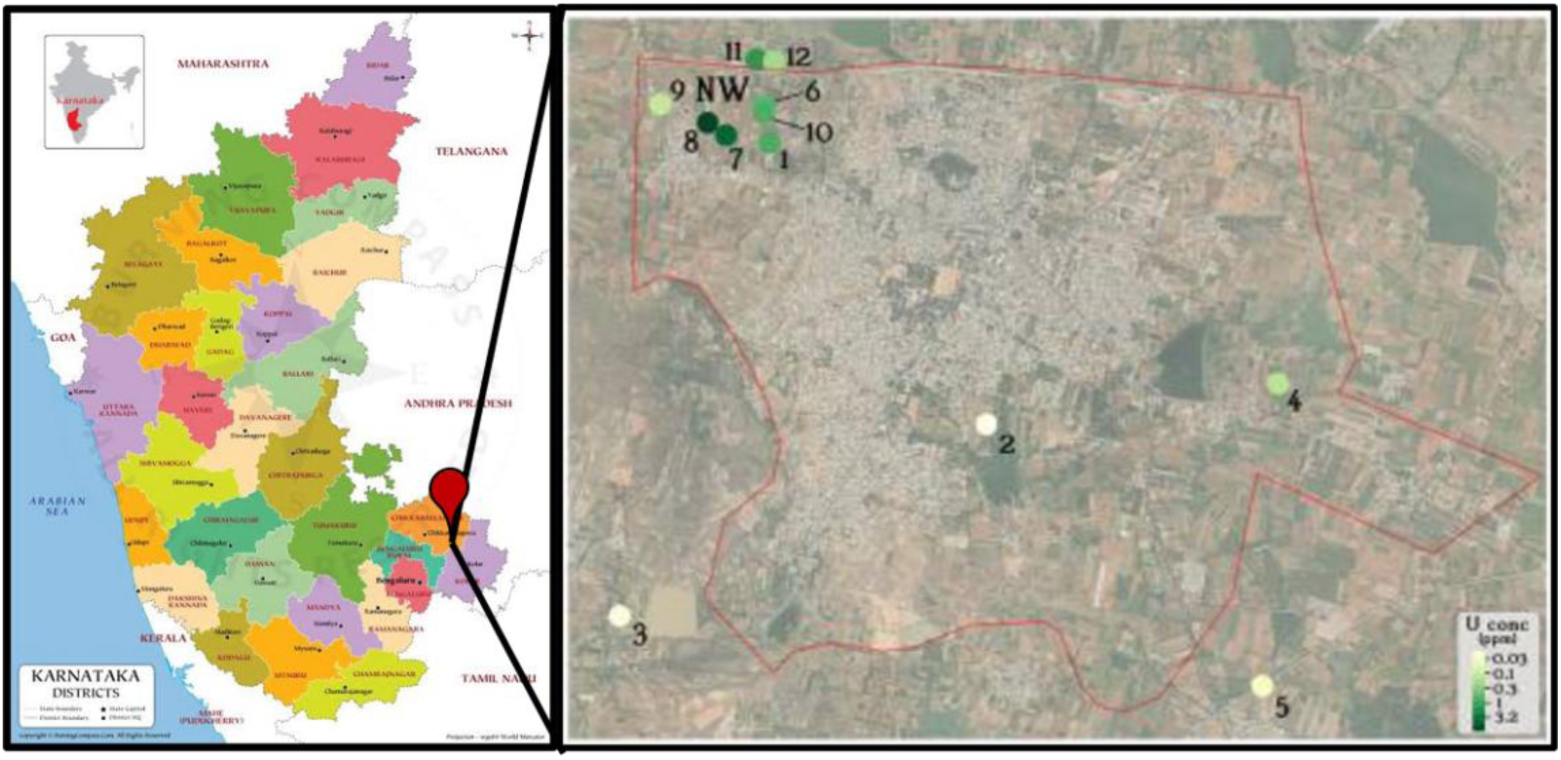
Map showing groundwater sample collection locations BW 1 to BW 12 (Details of the sample location are provided in our previous study [28].)

**Figure 2:**
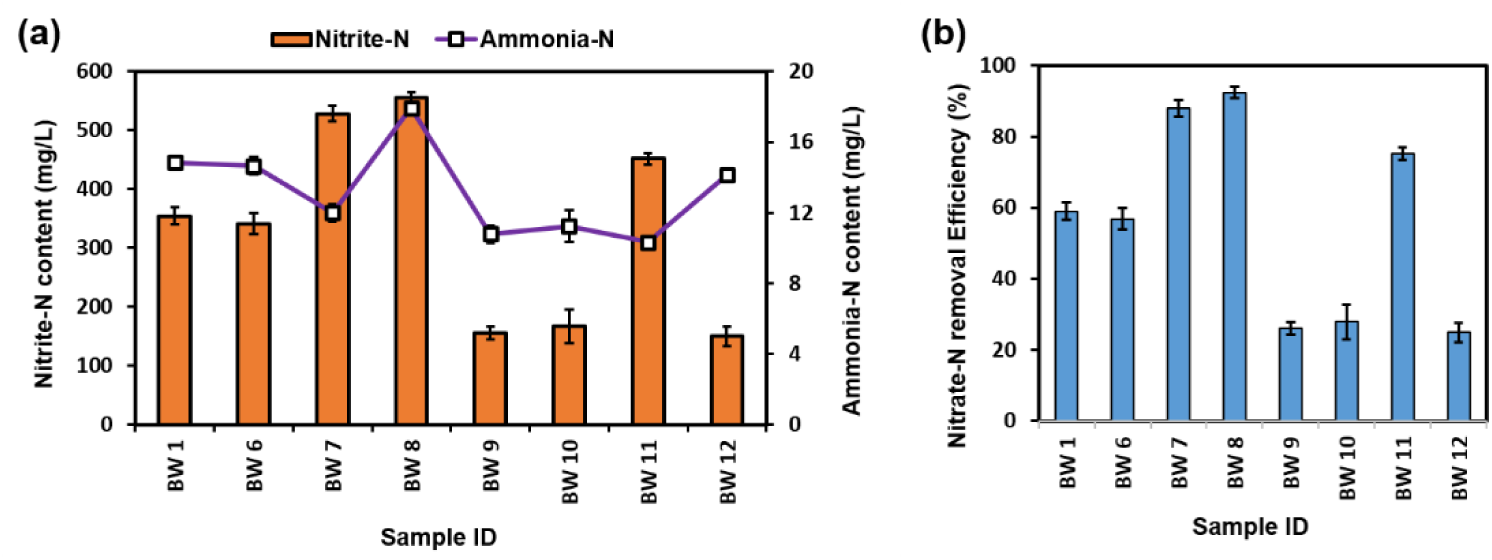
Screening of nitrate removal capacity of microbial consortia. (a) Quantitative estimation of Nitrite-N and Ammonium-N production by microbial consortia enriched from BW 1, BW 6, BW 7, BW 8, BW 9, BW 10, BW 11, and BW 12 water samples. (b) Nitrate removal efficiency after 10 hours of bacterial enrichment in MICP-Nitrate medium at 30°C.

Microbial consortia were enriched from BW 1, BW 6, BW 7, BW 8, BW 9, BW 10, BW 11, and BW 12 groundwater samples w.r.t. nitrate reduction and urea hydrolysis capabilities. The BW 2, BW 3, BW 4, and BW 5 groundwater samples are not part of this study due to negligible microbial growth recorded during enrichment study. The microbial consortia were allowed to grow in nitrate-reduction enrichment media (NB; nitrate broth), then allowed to induce under urea hydrolysis MICP selective media (NBUC) to facilitate step-by-step acclimatization of bacteria of interest from raw water samples [19].

### 2.3 Screening for Nitrite and Ammonia Production

Cell-free supernatants (CFS) from NB induction media was analyzed for nitrite production using the Griess colorimetric assay, and absorbance was measured at 540 nm. Ammonia production was determined using the Nessler’s colorimetric assay, and the absorbance was measured at 420 nm. Nitrite-N and ammonia-N concentrations in the CSF were directly proportional to the nitrate-N utilization by the microbial consortia. All analyses were performed in triplicate and all the data represented as mean value ± SD. The total nitrate removal efficiency was measured after 10 h of aerobic batch culture at 30°C with 100 rpm stirring conditions for all the BW groundwater samples.

### 2.4 Screening for biomineral precipitation

A biomass-biomineral precipitation estimation study was performed for the top 5 most efficient nitrate-removal consortia (BW1, BW6, BW7, BW8, and BW11). The biomass-biomineral precipitate from the NBUC induction media was collected by centrifugation at 5000 rpm for 10 min at room temperature. The wet weight of the biomass-biomineral precipitate was measured after 10 h of aerobic batch cultures, in the presence and absence of 100 ppb Pb^2+^.

### 2.5 Determination of Pb^2+^ toxicity and removal efficacy

Lead is a toxic compound; excessive Pb reduces bacterial viability and biomineral precipitation efficacy[15]. The effect of Pb^2+^ was studied only for BW 8. The study was performed by monitoring consortial growth and evaluating viable cells at hourly intervals. Microbial growth was recorded as O.D. at 600nm; a direct measure of cell scatter at 1 h of interval till 24 h. Lead toxicity was also studied with increasing Pb^2+^ content from 0-1000 ppb in NBUC medium. Microbial viability was recorded as relative turbidity of the culture. The bacterial growth at NBUC without Pb^2+^ (0 ppb) was considered as 100% viable. The cultures were incubated at 30 °C with shaking at 100 rpm for 24 hours. The CFS was subjected to acid digestion, and residual Pb^2+^ in the CFS was analysed using Inductively Coupled Plasma Optical Spectroscopy (ICP– MS). ICP-MS experiments to quantify lead concentration in the parts per mbllion (ppb) range were performed by Eurofins Scientific India using a PerkinElmer ^®^ 350X instrument. 40 mL of BW8 culture CFS from each experiemtal set were submitted for ICP-MS analysis. The instrument was set to detect elemental lead concentrations with an LOD of 0.01 ppb.

### 2.6 Identification and characterization of biomass-biomineral precipitate

Microbial consortia were identified using V3-V4 metagenomic analysis for BW 8-NBUC culture conditions. Total biomass was collected by centrifugation at 500 rpm for 10 min at room temperature. The microbial consortia analysis was outsourced to HealthVard Genomics (Health Vard Genomics Pvt. Ltd. Janai, Hooghly, India). The V3–V4 amplicon sequencing, targeting the V3 and V4 hypervariable regions of the bacterial 16S rRNA gene to identify and profile microbial communities.

The morphology and crystal structure of the biomineral precipitate were characterized using SEM, TEM-SEAD and XRD. All the samples were prepared from the total biomass-biomineral collected by centrifugation at 500 rpm for 10 min at RT. The precipitate was then fixed (2% w/v glutaraldehyde and 2% w/v paraformaldehyde) and dehydrated using graded ethanol (30% to 100%). SEM samples were gold-coated (Gold Sputtering Unit DESK II, Denton Vacuum, Moorestown, USA) for 120 s before imaging. The morphology of bacterial consortia and biomineral structures was imaged using FESEM (ThermoFisher® Apreo 2S HiVac). TEM-SEAD (ThermoFisher® Tecnai™ T20-ST) was performed by mounting the sample on the carbon-coated Cu-grid. Powdered samples were scanned by X-ray diffractometer (powder XRD, Rigaku Ultimate IV, Rigaku Corporation, Tokyo, Japan) and analysed for chemical components (Jade XRD pattern analysis software). The d-spacing and 2*θ* Peak values were compared with JCPDS / PDF Card No. 47-1734 (cerussite), 13-0131 (hydroxycerussite), and 05-0586 (calcite), matched from the International Centre for Diffraction Data (ICDD) search portal.

### 2.7 Experiemntal design and statistical analysis

The parametric optimization of a two-level factorial (2^4^) design was constructed using Design-Expert® software (Stat-Ease 360). The optimization experiment was performed in Nitrate Salt Medium (NSM) with BW8, selected as the highest biomass-biomineral precipitating microbial consortia, under NBUC culture conditions. The selected factors are a. initial nitrate-N, 2. Temperature, 3. pH, and 4. carbon concentration. Each factor was varied at two levels, namely low (-1) and high (+1), and the software-generated experimental matrix consisted of 16 runs. The details of the experimental design are summarized in Tables 2 and 3. Three responses were recorded for total nitrite-N production (NNP), total ammonia-N production (ANP), and total biomass-biomineral precipitation (BBP). The optimal condition was obtained with desirability function. In this design, importance +++ (3) was assigned to the selected responses. Analysis of variance (ANOVA) and other relevant statistical parameters generated by the software were used to evaluate the significance of the experimental factors and their effects on the measured responses.

**Table 2:** Factorial Design Parameters.

| Factor | Name | Units | Minimum | Maximum | Coded Low | Coded High |
| --- | --- | --- | --- | --- | --- | --- |
| A | Initial Nitrate-N | ppm | 100 | 600 | -1 | +1 |
| B | Temperature | °C | 23 | 37 | -1 | +1 |
| C | pH | - | 5 | 8 | -1 | +1 |
| D | C - conc. | g/L | 2.75 | 5.5 | -1 | +1 |

**Table 3:** Design matrix of optimization experiment and responses.

| Run | Factor 1<br>A: Initial Nitrate-N<br>(ppm) | Factor 2<br>B: Temperature<br>(°C) | Factor 3<br>C: pH<br>- | Factor 4<br>D: C - conc.<br>(g/L) |
| --- | --- | --- | --- | --- |
| 1 | 100 | 37 | 5 | 5.5 |
| 2 | 600 | 37 | 5 | 5.5 |
| 3 | 600 | 37 | 8 | 5.5 |
| 4 | 100 | 23 | 5 | 5.5 |
| 5 | 600 | 23 | 8 | 2.75 |
| 6 | 100 | 23 | 8 | 2.75 |
| 7 | 600 | 23 | 5 | 2.75 |
| 8 | 600 | 37 | 8 | 2.75 |
| 9 | 100 | 23 | 5 | 2.75 |
| 10 | 600 | 37 | 5 | 2.75 |
| 11 | 100 | 23 | 8 | 5.5 |
| 12 | 600 | 23 | 5 | 5.5 |
| 13 | 600 | 23 | 8 | 5.5 |
| 14 | 100 | 37 | 8 | 5.5 |
| 15 | 100 | 37 | 5 | 2.75 |
| 16 | 100 | 37 | 8 | 2.75 |

## 3. RESULTS AND DISCUSSION

### 3.1 Nitrate removal efficiency

The nitrate removal efficiency of microbial consortia from the borewell groundwater samples (BW 1, BW 6, BW 7, BW 8, BW 9, BW 10, BW 11, and BW 12) were screened by measuring the production of nitrite and ammonia in the batch culture. The consortia were predicted to participate in both denitrification and the Dissimilatory Nitrate Reduction to Ammonia (DNRA) process. Complete or partial denitrification involves the following reactions-

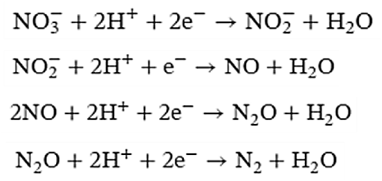

The Dissimilatory Nitrate Reduction to Ammonia (DNRA) process involves the following reactions-

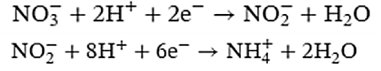

The production of ammonia confirms the DNRA process of nitrate reduction by all the microbial consortia. Total nitrite-N and ammonia-N production were measured and calculated against nitrite and ammonia standards. The top 5 samples were found to be BW8, BW7, BW11, BW1, and BW6, which produced > 300ppb of nitrite-N and >10 mg/L of dissolved ammonia-N (Figure 1a). The overall nitrate-N removal efficiency by microbial consortia of 92% for BW8, 88% for BW7, 75% for BW11 and around 50% for BW1 and BW6 was recorded (Figure 1b).

### 3.2 Pb^2+^ toxicity and Biomass-Biomineral precipitation

The biomass-biomineral wet weight was measured as an indicator of biogenic biomineralization with and without 100 ppb Pb^2+^ . The maximum biomass-biomineral of 15.38 ± 0.92 g/L was precipitated by BW 8 microbial consortia (Figure 3a). The further experiments involve only BW8 microbial consortia to explore microbial growth patterns and bacterial viability at elevated Pb^2+^ to evaluate toxicity. The standard sigmoidal growth pattern was obtained with and without 100 ppb Pb^2+^ , while about a 20% growth reduction was measured in presence of Pb^2+^ (Figure 3b). Figure 3c showing the specific growth rate (μ), which was calculated by fitting the exponential portion of the growth curve. Negligible effect was observed on the specific growth rate (μ= 0.707 h-1 without Pb^2+^ and μ= 0.679 h-1 with Pb^2+^), however, a slight elongated (+1h) lag phase was observed. Further, the effect of elevated Pb^2+^ to 1000 ppb was demonstrated by measuring viable cells using spread plate method. Taking no supplemented lead (0 ppb) as the 100% growth reference, viability was observed to be 84% for 100 ppb, 66% for 200 ppb, 43% for 400 ppb, 11% for 800 ppb and only 6% for 1000ppb (Figure 3d).

**Figure 3:**
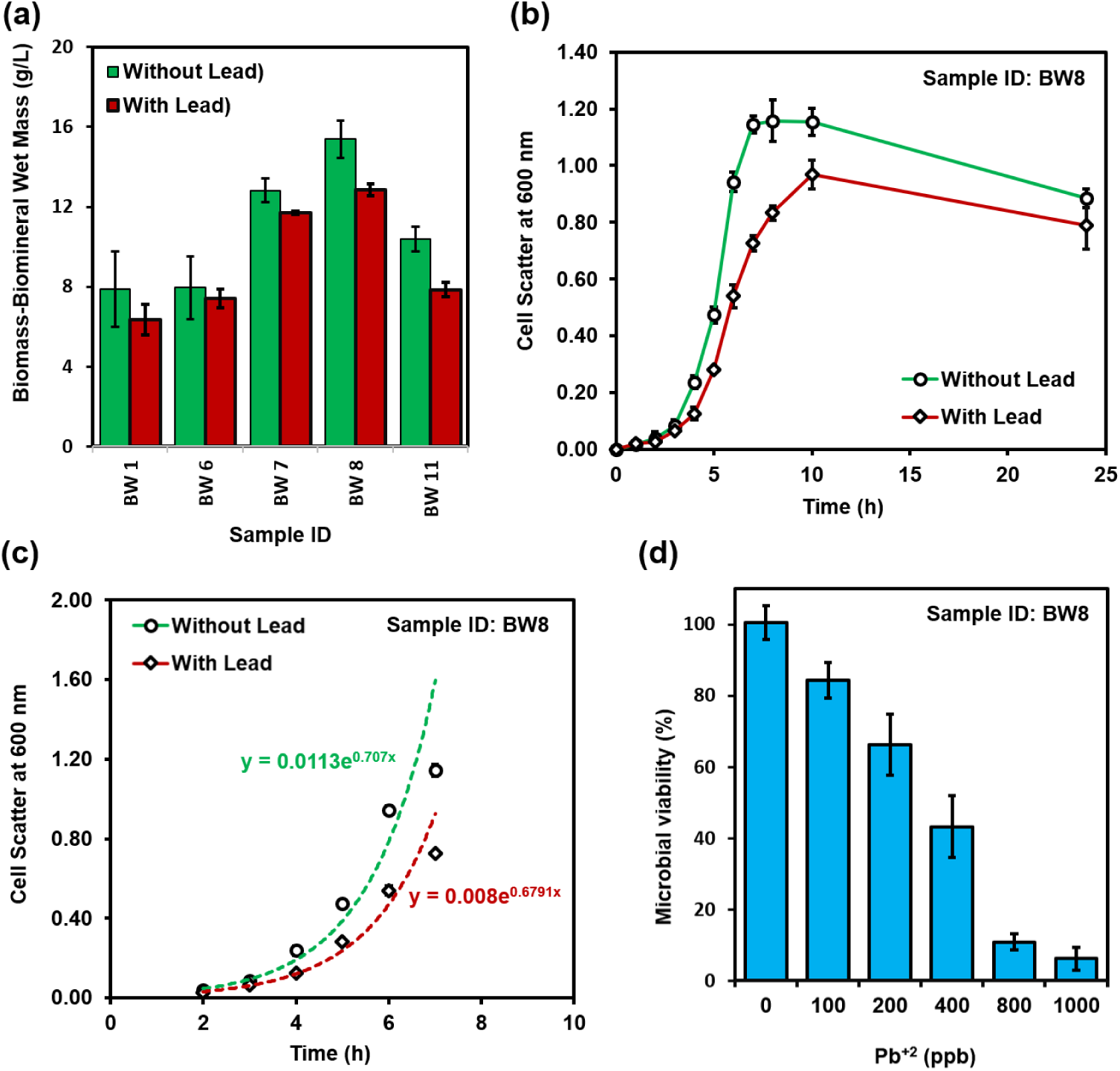
Screening of the biogenic mineralization capacity of microbial consortia (a) from BW 1, BW 6, BW 7, BW 8, and BW 11 water samples. (b) Microbial growth pattern, recorded with Lead (100 ppb) and without Lead, showing a substantial reduction in growth for the BW 8 water sample. (c) Exponential fit curve of the sigmoidal growth pattern of microbial consortia (BW 8), showing negligible differences in specific growth rate (μ) with Lead (100ppb Pb^2+^, μ = 0.679) and without Lead (μ = 0.707). (d) The microbial viability study at increased Pb^2+^ concentrations recorded after 10 hours of bacterial enrichment in MICP-Nitrate medium at 30°C. It confirms approximately 50% microbial viability at 400 ppb Pb^2+^, while showing a drastic reduction to about 10% at 800 ppb.

### 3.3 Biomass-Biomineral characterization

The biomass was characterized using 16S rRNA metagenomics for bacterial consortia identification and the biomineral was characterized using SEM, TEM-SEAD and XRD. The metataxonomic analysis using Krona plot of identified bacteria (Figure 4) shows the taxonomic distribution from phylum to species level and associated abundance based on the percentage of mapped reads considering NGS repositories based on the NCBI database. The microbial consortia was found to be 97% bacteria where about 40% bacteria belong to genus Bacillus and about 54% of bacteria belong to genus Acinetobacter.

**Figure 4:**
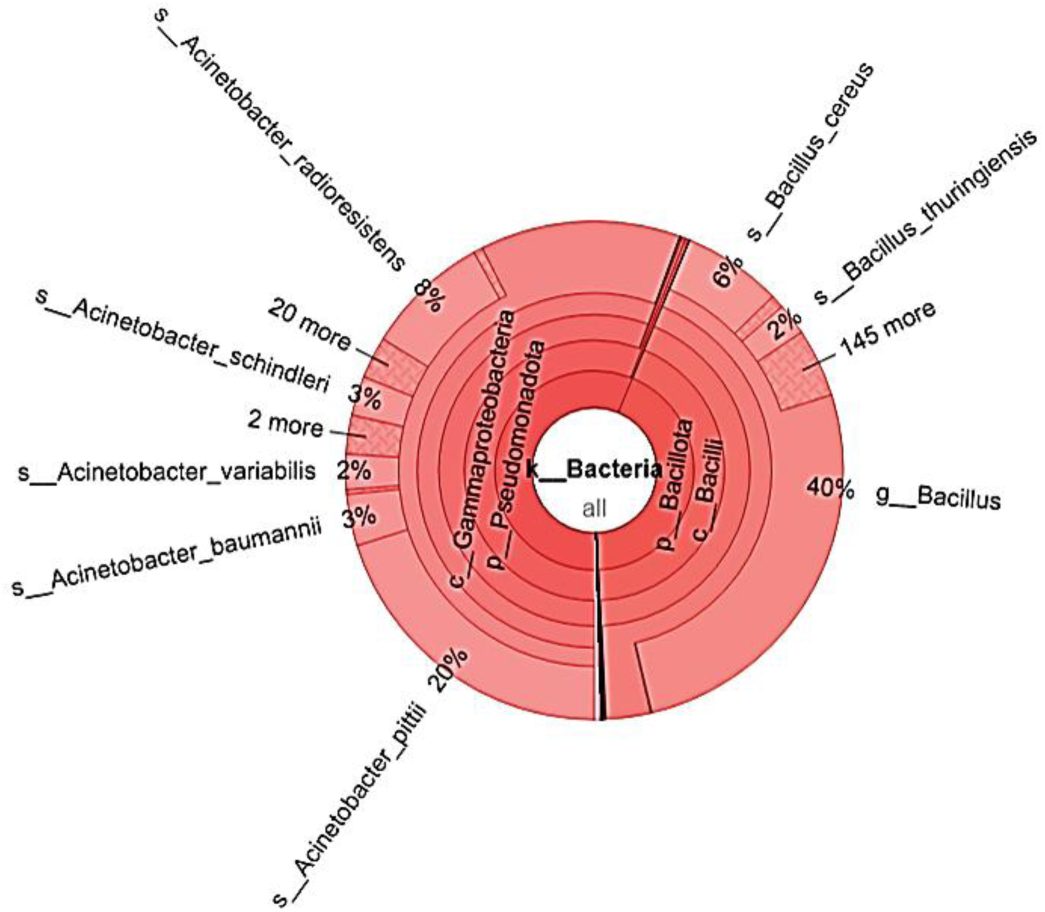
Metataxonomic Analysis of bacterial consortia identified from GW 8 induced in MICP Pb^2+^ precipitation conditions. Among the 97% of total bacteria, 39% belong to the genus Bacillus, and 54% belong to the genus Acinetobacter.

The SEM image of bacterial consortia along with biomineral precipitated via MICP showed (Figure 5a) granular deposition on and around the bacterial cells throughout the samples. The same was visualized under TEM at enlarged single bacteria level to confirm the location of biomineral formation. TEM images reconfirmed the nano-sized biomineral precipitation (Figure 5b). The selected area diffraction (SEAD) showed the crystalline nature of the biomineral (Figure 5b, inset) where the d-spacings of 3.54, 2.83, 1.88, 1.52, and 1.14 Angstrom calculated by fitting the ring using ImageJ (version 1.54g), matched the lead carbonate JCPDS card. Additionally, powder XRD results confirmed the precipitation of calcite biomineral in the absence of Pb^2+^, whereas co-precipitation of calcite (CaCO_3_) and cerussite (PbCO_3_) was observed in the presence of Pb^2+^ (Figure 5c). The XRD peaks at 2*θ* positions (104), (110), (113), (202), (016), and (018) confirm calcite formation. On the other hand, typical XRD peaks at the 2 theta positions (111), (200), (112), (202), and (130), along with the calcite peaks, confirm the co-precipitation of cerussite with calcite. An enlarged biomineral particle was also explored using TEM-SAD (Figure 5d) and analyzed for d-spacing measurement using ImageJ software (version 1.54g). HR-TEM of the enlarged biomineral particle displayed a calculated d-spacing of 2.66 nm.

**Figure 5:**
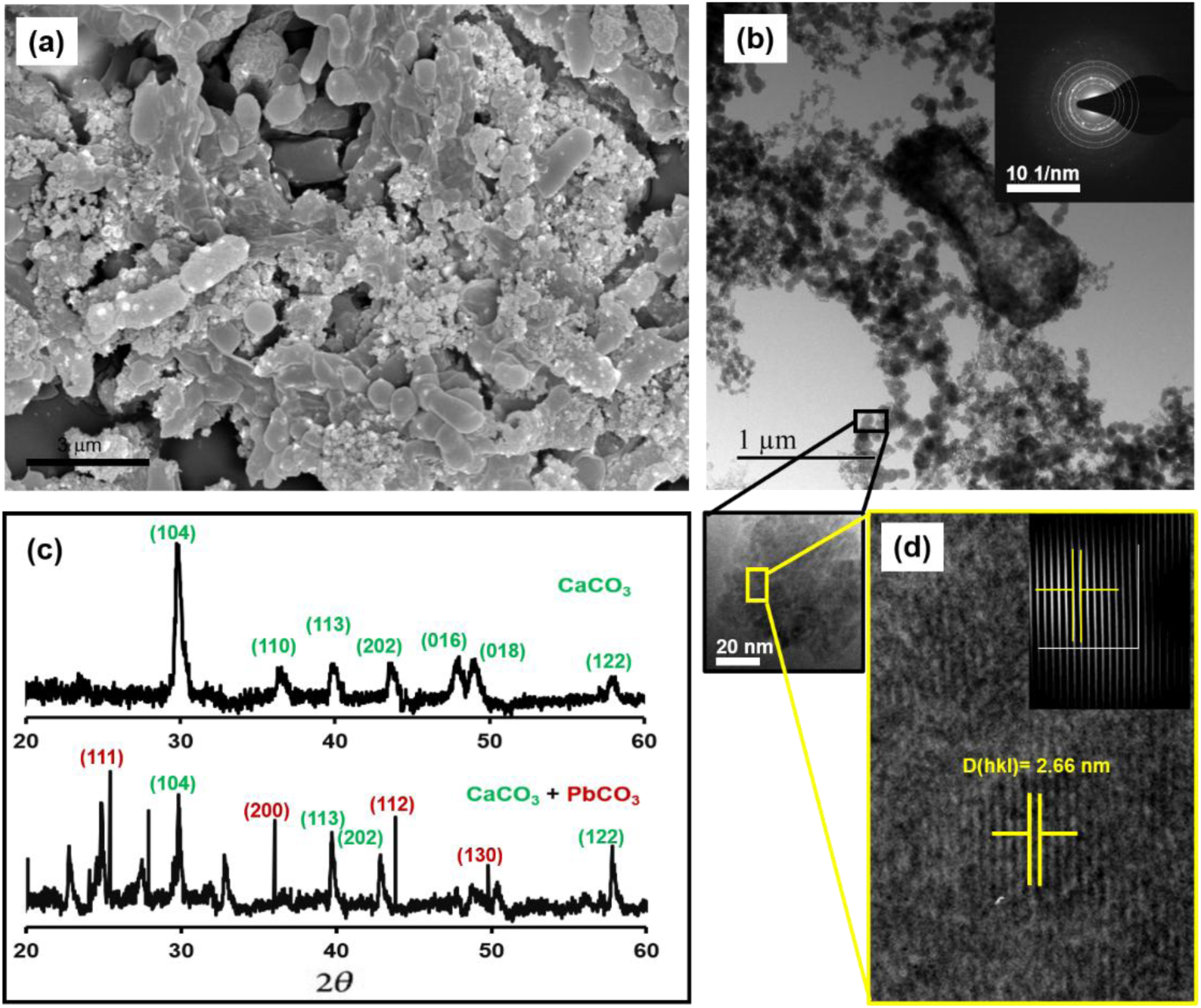
Characterization of biogenic minerals precipitated in GW 8. (a) Scanning electron microscopy (SEM) image showing bacterial consortia with granular precipitation on the bacterial cell surfaces. (b) Transmission electron microscopy (TEM) image with selected area diffraction pattern (SEAD, inset) showing biomineral precipitation on and around the bacterial cell, with d-spacings of 3.54, 2.83, 1.88, 1.52, and 1.14 Angstrom calculated by fitting the ring using ImageJ (version 1.54g), matched with the lead carbonate JCPDS card. (c) XRD intensity plot of biomass-biomineral obtained with and without Pb^2+^ supplement. Miller indices clearly indicate the co-precipitation of PbCO_3_ with CaCO_3_, indicating the formation of calcite and cerussite in the induction media. (d) HR-TEM of the enlarged biomineral particle displaying a calculated d-spacing of 2.66 nm. Analysis was performed using ImageJ software (version 1.54g). (d)-Inset showing inverse FFT image of selected area of interest.

### 3.4 Optimization model

A total of 16 experiments were designed for biomineral precipitation using two-level (2^4^) full factorial methodologies. The model evaluation suggested linear model with no aliased factors for all the responses. The linear model showed high adjusted R^2^, predicted R^2^ and calculated *p*-values. High adjusted R^2^ and low *p*-value indicated the data fit the model well, while high predicted R^2^ suggested the model could provide good estimation of new responses. The Predicted R² of 0.7663 was in reasonable agreement with the Adjusted R² of 0.9041, while the calculated difference is less than 0.2. Adequate precision requires a signal-to-noise ratio greater than 4. The present model’s ratio of 17.636 indicates an adequate signal, which also suggests that this model can be used to navigate the design space. The model equations were predicted and constructed for the nitrite-N production, ammonia-N production, and biomass-biomineral mass precipitation using Design-Expert®. The experimental responses are shown in Table 4. Table 5 shows the numerical model that predicts all three responses in terms of coded and actual values of the factors. These models are correlational functions that can be used to estimate the responses from known variables.

**Table 4:** Responses recorded in the design matrix.

| Run | Response 1<br>NNP<br>(ppb) | Response 2<br>ANP<br>(ppm) | Response 3<br>BBP<br>(ppb/mg/L) |
| --- | --- | --- | --- |
| 1 | 93 | 77 | 16.9 |
| 2 | 112 | 72 | 16.43 |
| 3 | 648 | 73 | 28.67 |
| 4 | 115 | 61 | 13.4 |
| 5 | 622 | 59 | 24.2 |
| 6 | 108 | 59 | 13.77 |
| 7 | 93 | 66 | 20.03 |
| 8 | 140 | 97 | 20.43 |
| 9 | 86 | 49 | 16.43 |
| 10 | 49 | 74 | 19.9 |
| 11 | 117 | 33 | 18.1 |
| 12 | 64 | 34 | 14.17 |
| 13 | 220 | 37 | 18.87 |
| 14 | 107 | 54 | 18.87 |
| 15 | 73 | 48 | 17.6 |
| 16 | 131 | 89 | 18.87 |

**Table 5:** Model equations.

| Responses | Coded values |
| --- | --- |
| NNP | $= 173.62 + 69.87 A - 4.50B + 88.00 C + 10.87 D + 76.00$<br>$AC + 60.00 BD + +65.25 ABD + 49.63 BCD + +52.62 ABCD$ |
| ANP | = 61.38 +2.62 A +11.62 B -6.25 D -7.13 CD |
| BBP | =18.54 +1.80 A +1.17 B +1.68 C +1.02 AC +0.8725 BD +1.27 CD +1.12 ABD +1.12 ABCD |

**Table 6:**
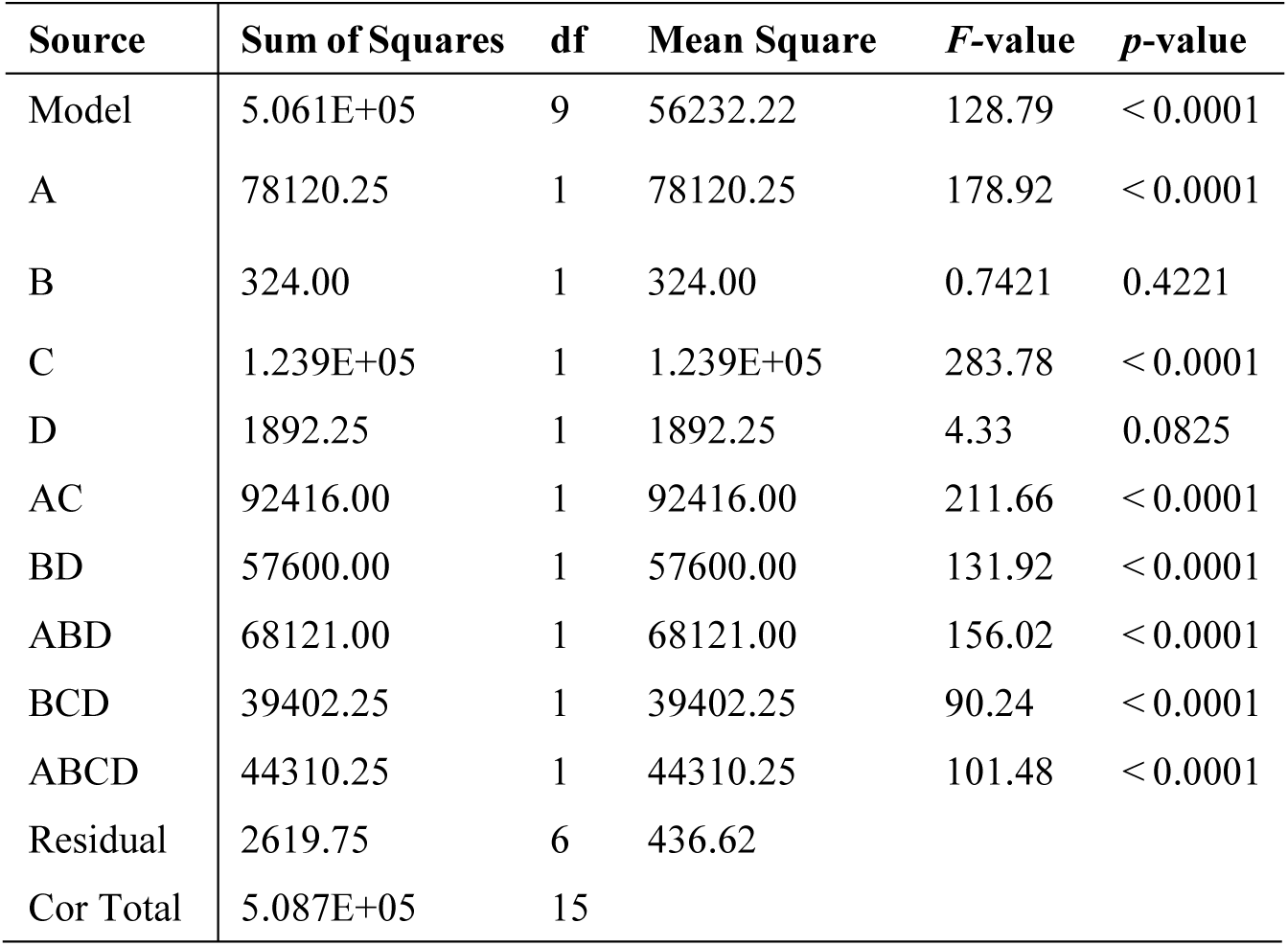
ANOVA analysis using coded values for nitrite-N production (NNP).

| Source | Sum of Squares | df | Mean Square | F-value | p-value |
| --- | --- | --- | --- | --- | --- |
| Model | 5.061E+05 | 9 | 56232.22 | 128.79 | < 0.0001 |
| A | 78120.25 | 1 | 78120.25 | 178.92 | < 0.0001 |
| B | 324.00 | 1 | 324.00 | 0.7421 | 0.4221 |
| C | 1.239E+05 | 1 | 1.239E+05 | 283.78 | < 0.0001 |
| D | 1892.25 | 1 | 1892.25 | 4.33 | 0.0825 |
| AC | 92416.00 | 1 | 92416.00 | 211.66 | < 0.0001 |
| BD | 57600.00 | 1 | 57600.00 | 131.92 | < 0.0001 |
| ABD | 68121.00 | 1 | 68121.00 | 156.02 | < 0.0001 |
| BCD | 39402.25 | 1 | 39402.25 | 90.24 | < 0.0001 |
| ABCD | 44310.25 | 1 | 44310.25 | 101.48 | < 0.0001 |
| Residual | 2619.75 | 6 | 436.62 |  |  |

|  |  |  |
| --- | --- | --- |
| Cor Total | 5.087E+05 | 15 |

**Table 7:**
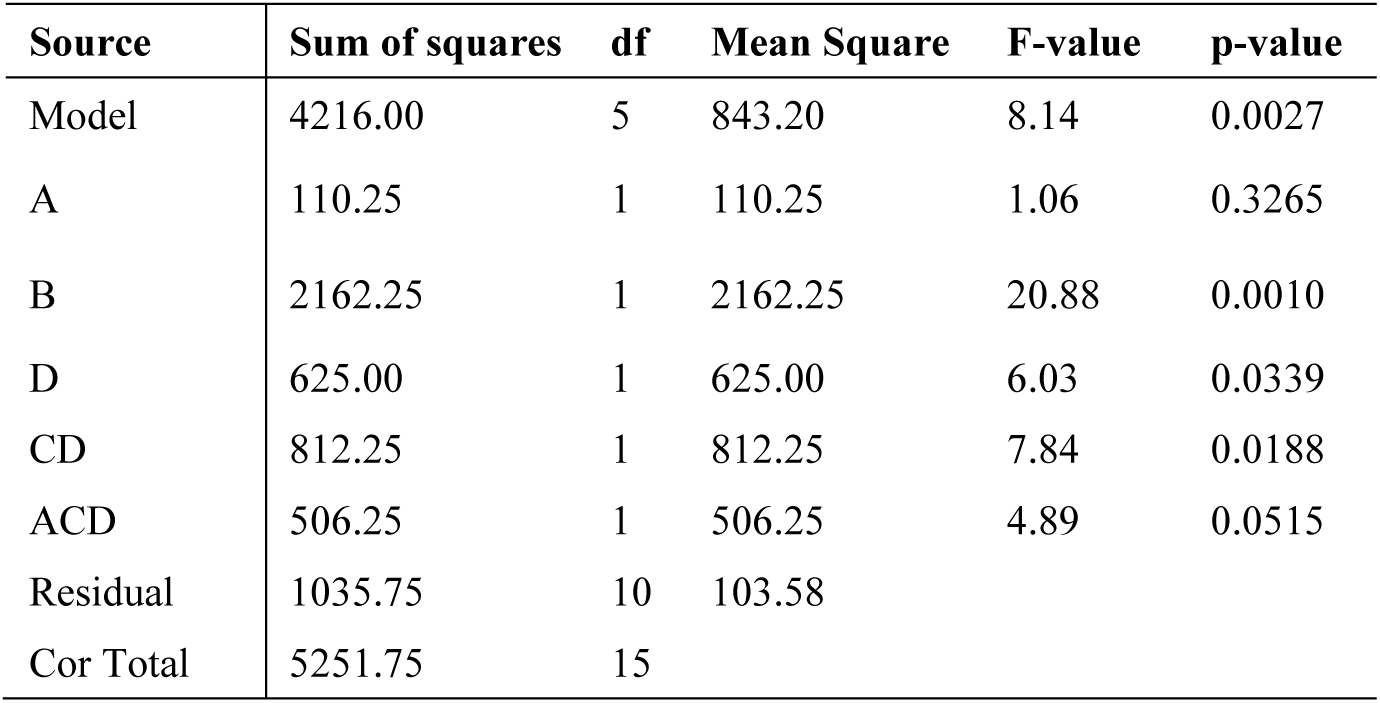
ANOVA analysis using coded values for ammonia-N production (ANP).

**Table 8:** ANOVA analysis using coded values for biomass-biomineral precipitation (BBP).

| Source | Sum of Squares | df | Mean Square | F-value | p-value |
| --- | --- | --- | --- | --- | --- |
| Model | 213.56 | 8 | 26.70 | 18.68 | 0.0005 |
| A | 51.70 | 1 | 51.70 | 36.18 | 0.0005 |
| B | 21.86 | 1 | 21.86 | 15.30 | 0.0058 |
| C | 45.29 | 1 | 45.29 | 31.70 | 0.0008 |
| AC | 16.73 | 1 | 16.73 | 11.71 | 0.0111 |
| BD | 12.18 | 1 | 12.18 | 8.53 | 0.0223 |
| CD | 25.76 | 1 | 25.76 | 18.03 | 0.0038 |
| ABD | 20.16 | 1 | 20.16 | 14.11 | 0.0071 |
| ABCD | 19.89 | 1 | 19.89 | 13.92 | 0.0073 |
| Residual | 10.00 | 7 | 1.43 |  |  |
| Cor Total | 223.56 | 15 |  |  |  |

Analysis of variance (ANOVA) of the developed models was performed and the results are shown in Tables 6 (for NNP) 7 (for ANP), and 8 (for BBP). The ANOVA test was usedto evaluate the model fitting to the responses with a confidence level of 95%. All three models were significant (*p*-value < 0.01). The adequate precisions were greater than 4, demonstrating that the signal-to-noise ratio was sufficient to detect the effect of each factor. The initial nitrate (A) concentration showed a significant effect (p-value<0.01) on the nitrite production and biomass-biomieral precipitaion; however, it did not show a significant effect on ammonia production (*p*-value *=* 0.32). The initial pH of the culture medium showed a significant effect as well. On the other hand, the main effect of temperature was insignificant, while the interaction effect with multiple factors showed a significant effect on NNP. Similarly, pH didn’t show a significant effect as a main factor; however, in combination with other factors (AC, BCD, and ABCD; *p-*value < 0.1) on ammonia production. For the biomass-biomineral precipitation, the added carbon content (factor D) did not show a significant effect, whereas other factors significantly affected the biomineral precipitation. These observations suggest that the carbon source in the culture medium is not the carbon source of the carbonate precipitated in the medium.

The validity of the model was analyzed using predicted vs. actual values plots (Figure 6) and residual vs. predicted value plots (Figure 7) for all three responses. No significant outliers were observed, and no systematic trend was present for the residuals. This suggested that the model fit is accurate.

**Figure 6:**
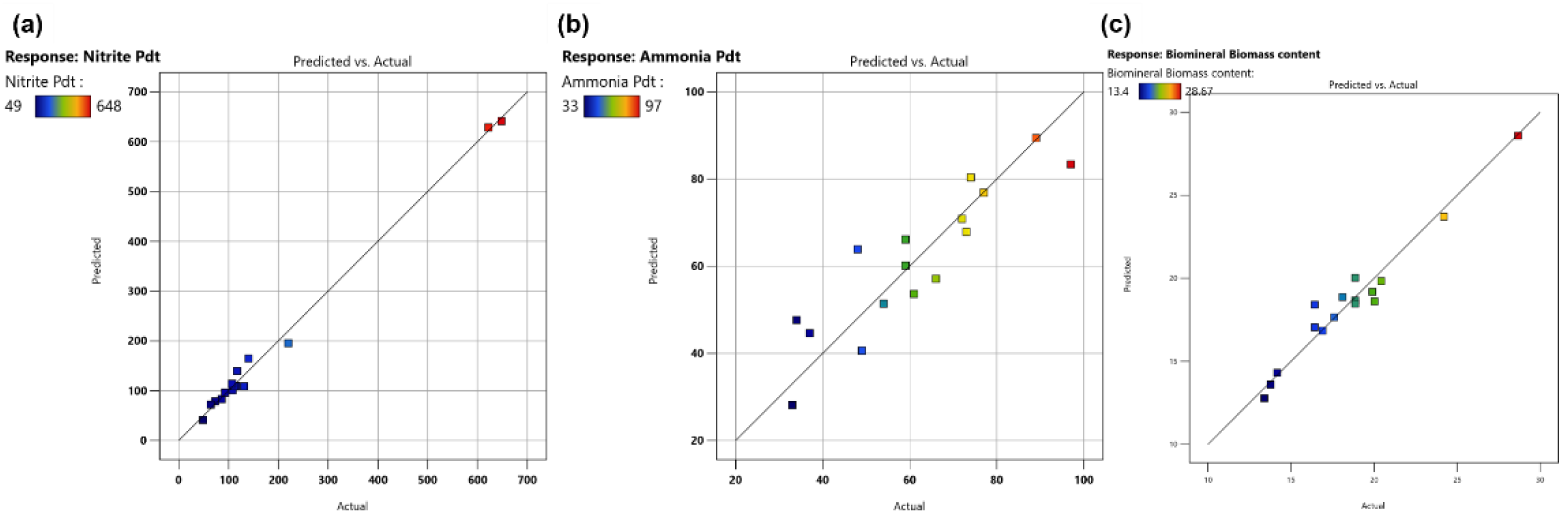
Predicted vs. actual data for NNP (a), ANP (b) and BBP (c).

**Figure 7:**
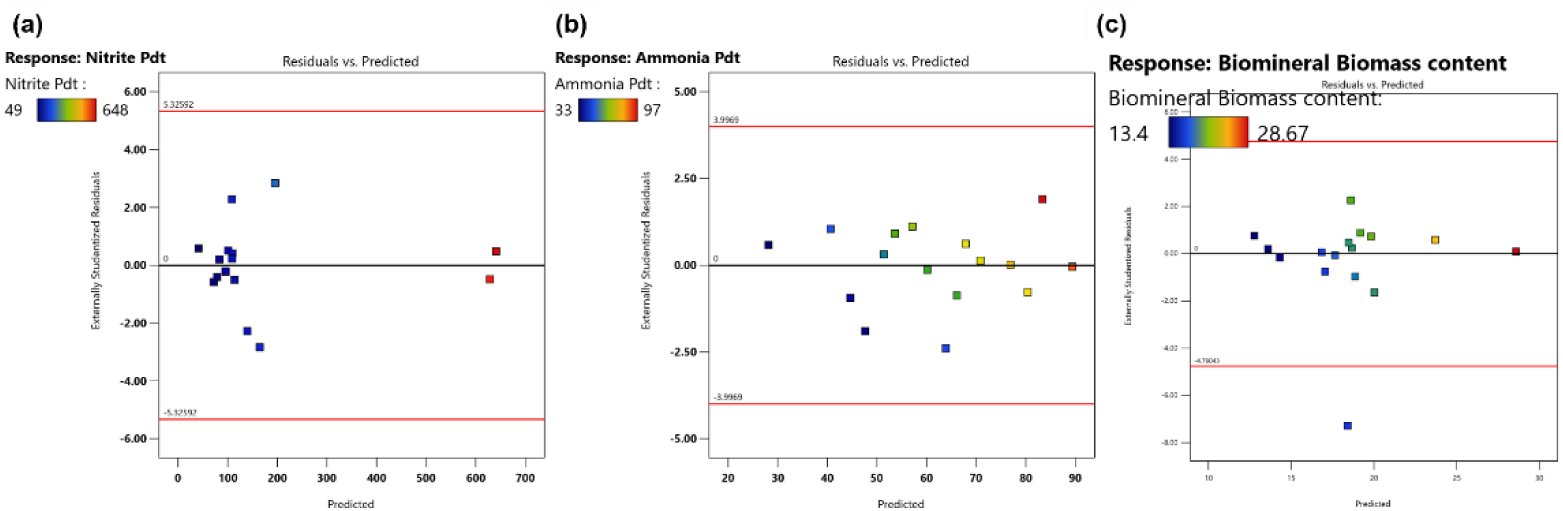
Residual vs. Predicted data for NNP (a), ANP (b) and BBP (c).

The main effects were found to be very straight, with increasing initial nitrate resulting in higher nitrite production, higher pH elevating ammonia production, thus facilitating increased biomass-biomineral precipitation. The interacting factors showed interesting effects. Contour plots were employed to explore the interacting factors for all three responses in Figure 8. The red dots are the experimental data points displayed on the plots. Figures 8a and 8b show the interaction between initial nitrate and carbon source added to the culture media. A differential trend was observed when the pH was maintained at 8, while the culture was maintained at low (23 °C for Fig. 8a) and high (37 °C for Fig. 8b) temperatures. At lower temperatures, the lower carbon availability (2.75 g/L) with higher initial nitrate (600 ppm) was predicted to produce higher nitrite. Inversely at higher temperatures, the higher carbon availability (5.5 g/L) with higher initial nitrate (600 ppm) was predicted to produce higher nitrite. Comparatively, a little effect was predicted in case of ammonia production with culture temperatures (Figure 8c and 8d) throughout the carbon and nitrate content gradient. The factorial effect on the BBP showed a similar trend for all three factors (Figure 9). When pH and C-content were kept at higher values, biomineral precipitation was predicted high towards high temperature and high nitrate level (Figure 9a). When temperature and C-content were kept at higher values, biomineral precipitation was predicted high towards high pH and high nitrate level (Figure 9b). When temperature and pH were kept at higher values, biomineral precipitation was predicted high towards high C-content and high nitrate level (Figure 9c).

**Figure 8:**
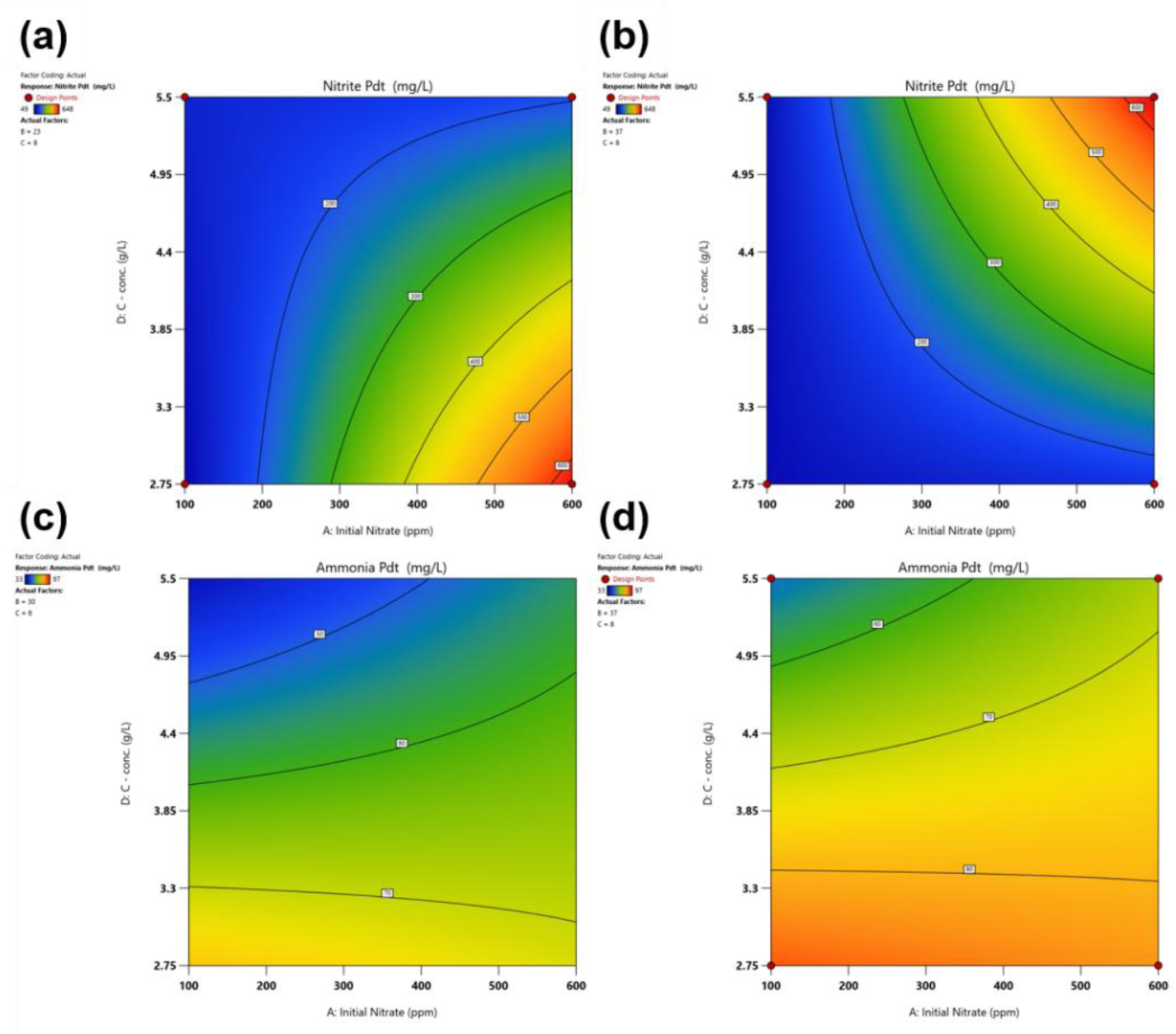
Contour plots of interactive effects of the four factors on (a) NNP and (b) ANP.

**Figure 9:**
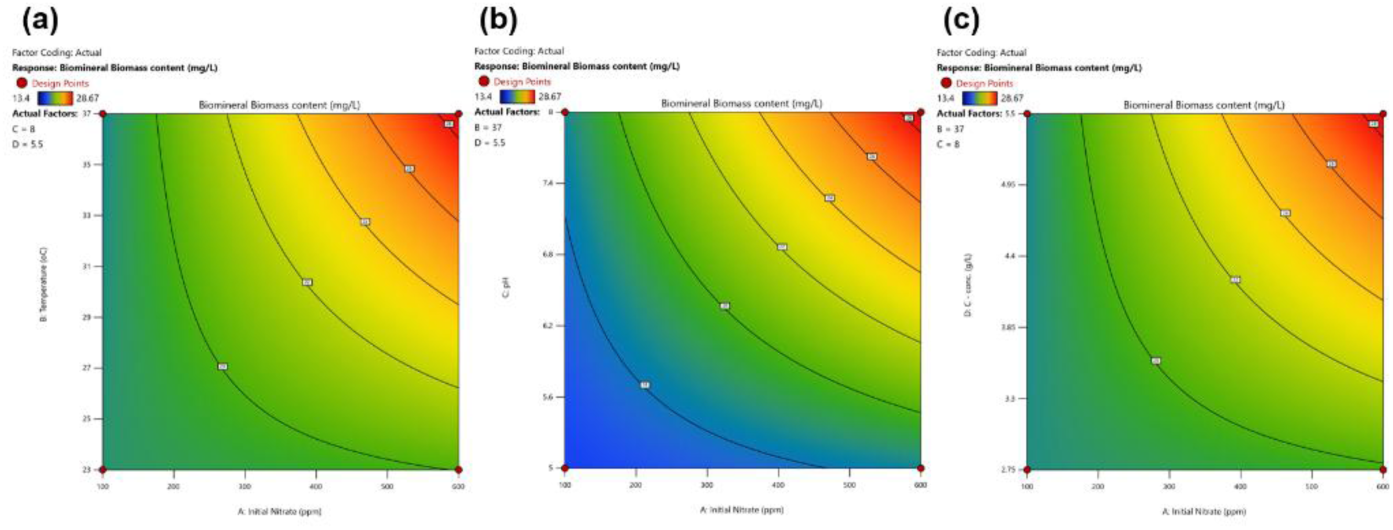
Contour plots of interactive effects of the three factors (a) temperature, (b) pH, and (c) carbon content w.r.t. initial nitrite on BBP.

Using thepredictive models, optimization of the nitrate reduction mediated Pb biomineralization conditions was conducted in Design Expert®, resulting in the initial nitrate requirement of 600 ppm, temperature 37 °C, pH 8, and carbon content of 5.5 g/L (Table 9). Confirmatory experiments were conducted in triplicates using the predicted points by the model. The predicted responses were NNP of 160 ± 26 ppb, ANP of 16 ± 4 ppm, and BBP of 17 ± 0.9ppb/mg/L . The predicted values were close to experimental data, showing that the models are relatively accurate (Table 10).

**Table 9:**
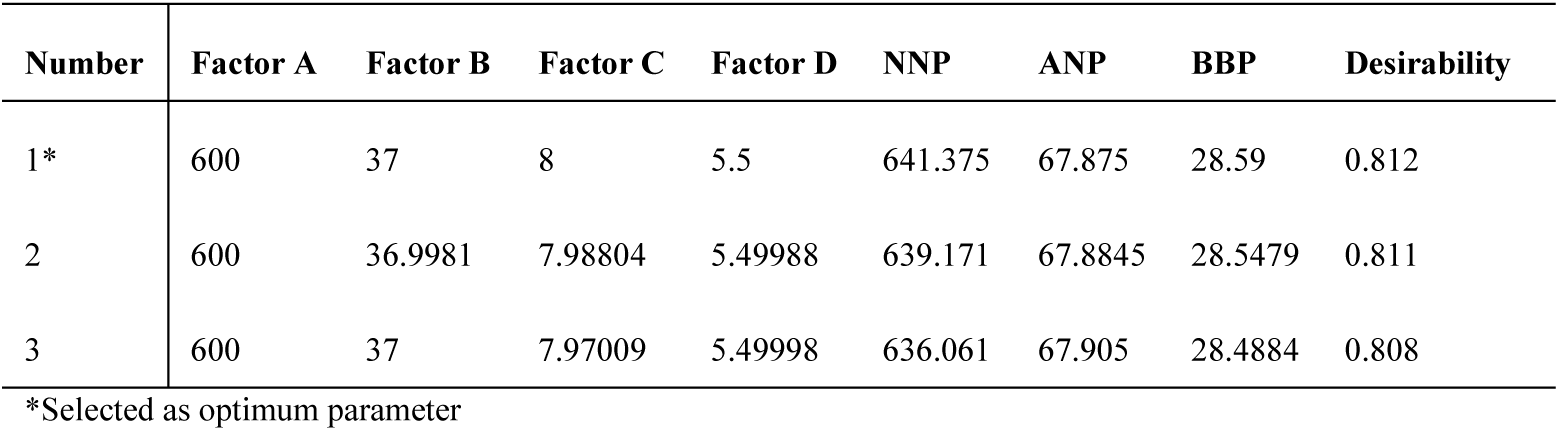
Optimal biogenic Pb remediation conditions.

**Table 10.**
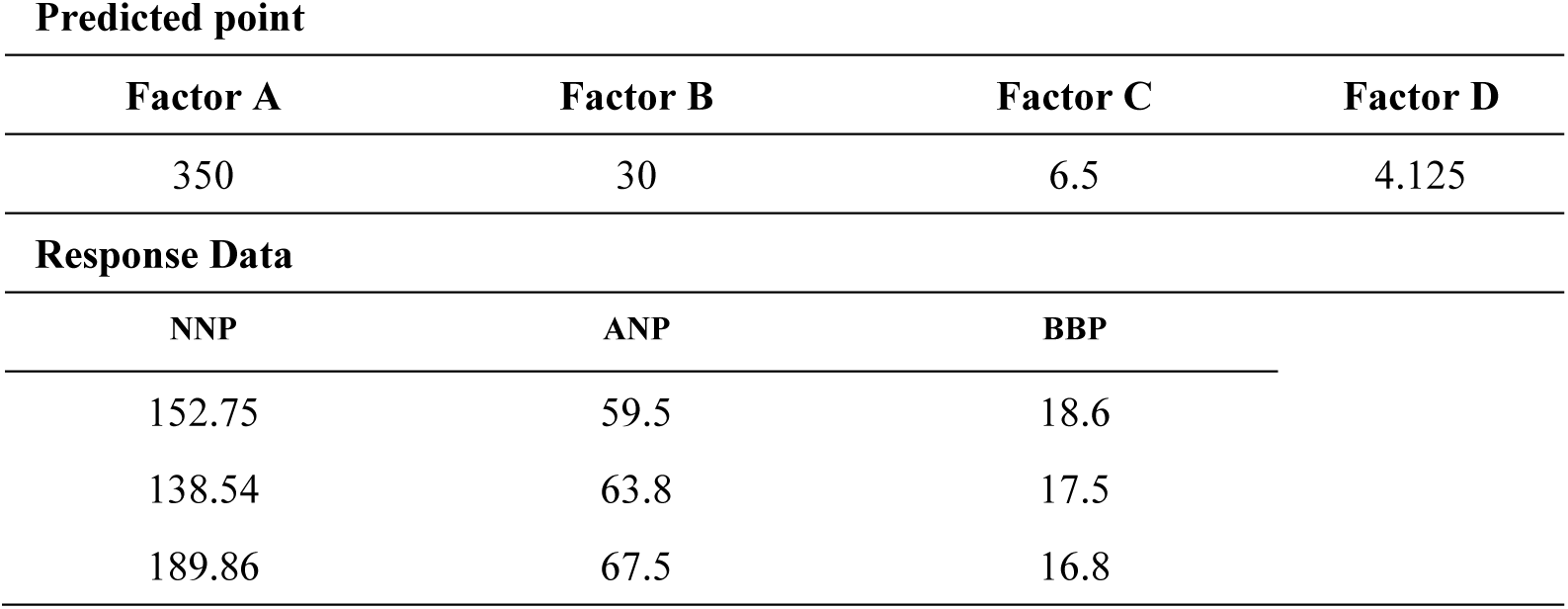
Confirmatory factor values and response data at the predicted point.

## 4. CONCLUSION

The present study demonstrates that indigenous bacterial consortia isolated from contaminated groundwater in Eastern Karnataka (Chintamani Village) showed the potential to simultaneously promote nitrate reduction, ammonia production, and microbially induced calcite precipitation (MICP) mediated biogenic lead mineralization. The selected bacterial consortia exhibited sustained nitrate reduction, nitrite accumulation, ammonia production and progressive alkalinization, confirming the combined activities of nitrate reductase and urease under the investigated conditions. Optimisation using Design-Expert® software identified favourable operating conditions for microbial growth and nitrogen transformation, while XRD and TEM-SEAD analysis confirmed the formation of biogenic minerals containing calcium, and lead, indicating the potential incorporation or co-precipitation of lead carbonate and calcium carbonate within the biomineralized matrix. This study provides a clear view on the parameters to be set for pilot-scale translation for a bioremedial solution towards Pb^+2^ contamination in groundwater.

## 6. Acknowledgements

TG coined the research idea, AB and SH performed the study. The research was funded by the Department of Biotechnology, Govt. of India, under the research grant of DBT-Ramalingaswami Fellowship awarded to TG. All the authors would like to thankfully acknowledge the funding and facilities provided by Ramaiah University of Applied Sciences, Department of Biotechnology, Bangalore. TG also gratefully acknowledges DN for water quality and physicochemical data of the samples used in this research.

